# Effects of Spin4 ablation in aging mice: body composition, bone density, and malignancy prevalence

**DOI:** 10.64898/2026.02.05.704000

**Authors:** Julian C. Lui, Isabelle Hannula, Arun Rama-Krishnan, Lijin Dong, Jeffrey Baron

**Affiliations:** Section on Growth and Development, Eunice Kennedy Shriver National Institute of Child Health and Human Development, National Institute of Health, Bethesda, Maryland, USA; Genetic Engineering Core, National Eye Institute, National Institute of Health, USA

**Keywords:** Overgrowth, Spindlin, Tumorigenesis, histone reader, epigenetics

## Abstract

Many overgrowth syndromes are associated with increased risk of tumorigenesis and malignancies. Our group recently identified a frameshift variant in histone reader *SPIN4* located on the X chromosome to be a new genetic cause for human overgrowth. In the current study, we investigated the prevalence of malignancies, along with body weight, body length, body composition and bone mineral density, in Spin4 knockout mice at 18 months of age. We found that male mice lacking Spin4 have increased number of tumors and increased body length, while body weight, body composition and bone mineral density were comparable with wild-type mice. We also analyzed publicly available expression data in various types of human cancers and looked for increased or decreased expression of genes that are implicated in overgrowth syndromes and act through epigenetic mechanisms. We found that the expression of *SPIN4, EZH2*, and *DNMT3A* to be elevated in many human cancers compared to the corresponding non-malignant tissue samples. Taken together, our current findings confirm that loss of SPIN4 causes overgrowth in mice (in terms of body length) and is associated with increased prevalence in neoplasia; but does not appear to affect adiposity or bone density.

## Introduction

Overgrowth syndromes represent a heterogeneous group of clinical conditions characterized by excessive prenatal or postnatal growth, which can be generalized or segmental (1). Overgrowth syndromes are frequently associated with increased risk of malignancies (2), many of which occur during childhood (3). For example, both children with Beckwith– Wiedemann syndrome (BWS) and Simpson-Golabi-Behmel syndrome (SGBS) are more likely to develop hepatoblastoma and Wilms tumor (4, 5). Patients with Sotos syndrome show an increased prevalence of neuroblastoma and teratoma (6). Similarly, Weaver syndrome is associated with higher risks for Hodgkin disease, acute lymphoid leukemia, and neuroblastoma (7).

Our group recently reported a male individual with an X-linked generalized overgrowth syndrome of prenatal onset (OMIM #301114) (8). Genetic study of the family identified a frameshift variant in *SPIN4*, which is located on the X chromosome and encodes histone reader protein Spindlin-4. More recently, a second family with two cases of overgrowth was identified to be associated with SPIN4 frameshift mutations, confirming the causal relationship between SPIN4 and human overgrowth (9). Together with several other genes such as *NSD1* (Sotos syndrome (10)), *EZH2* (Weaver syndrome (11)), *DNMT3A* (Tatton-Brown-Rahman syndrome (12)) and *CHD8* (CHD8-neurodevelopmental disorder with overgrowth (13)), *SPIN4* became the latest addition to a small group of epigenetic readers/writers/erasers that cause human overgrowth syndromes.

The long-term effects of Spin4 during aging have not been studied. Given the previously reported association between cancer risk and overgrowth syndromes, we hypothesized that patients with SPIN4 mutations may also have increased prevalence for malignancies. However, to date only a very small number of patients with SPIN4 mutations have been identified, making any clinical study on cancer risk of SPIN4 patients impractical. In the current study, we sought to examine this hypothesis in animal model by ascertaining the prevalence of neoplasms in mice lacking Spin4 at 18 months of age. We also analyzed publicly available expression data in different types of human cancers and looked for increased or decreased expression of genes that are implicated in overgrowth syndromes and act through epigenetic mechanisms, including SPIN4. We also evaluated the effects of Spin4 ablation on body composition and on bone mineral density because of the clinical importance of these endpoints in human aging.

## Materials and Methods

### Animal procedures

All animals were used in accordance with the Guide for the Care and Use of Laboratory Animals (National Research Council 2003). We previous generated C57BL/J mice carrying truncating mutations in Spin4 (1). Female heterozygous Spin4 (Spin4^+/-^) mice were mated with wild-type male mice to produce male or female wild-type mice, male hemizygous (Spin4^Y/-^) mice, or female heterozygous (Spin4^+/-^) mice for study. This mating scheme does not produce female homozygous (Spin4^-/-^) mice. Spin4^Y/-^ male mice, Spin4^+/-^ female mice, and wild-type male and female littermates were fed ad lib until 18 months of age, when they were sacrificed by carbon dioxide inhalation (30% flow rate of chamber volume per minute). Body weight and length measurements were taken, followed by body composition and bone mineral density measurement using the Xpert 80-L Specimen Radiography System (KUBTEC Scientific). Necroscopy was performed by veterinary pathologists in the Division of Veterinary Resources, National Institutes of Health (NIH). Tissue masses that were suspicious for a neoplasm on gross examination were examined further by histopathology.

### Bioinformatic analysis

Publicly available RNA-Seq data from 26 different cancer types along with matched normal samples was obtained from The Cancer Genome Atlas Program (TCGA) and downloaded using the Genomic Data Commons Data Portal (https://portal.gdc.cancer.gov/). Expression data for our genes of interest, which include epigenetic readers/writers/erasers that cause overgrowth syndromes (SPIN4, NSD1, EZH2, CHD8), an oncogene (CCND1), and a tumor suppressor gene (RASGRP2), were extracted from the dataset for comparison and data visualization.

## Results

### Increased body length in adult Spin4^Y/-^ mice

Hemizygous Spin4^Y/-^ male mice, heterozygous Spin4^+/-^ female mice, and their wild-type male and female littermates were sacrificed at 18 months of age, and their body length and weight were measured (**Figure 1**). We found that, in male mice, body length was significantly increased in Spin4^Y/-^ compared to wild-type littermates (**Figure 1A**, 10.80 cm vs 10.28 cm, P=0.002), while in females, Spin4 heterozygous only showed a tendency toward increased body length without reaching statistical significance (10.31cm vs 10.01cm, P=0.1281). In both male Spin4^Y/-^ and female Spin4^+/-^, mean body weight was slightly greater than in wild-type mice without reaching statistical significance (**Figure 1B**, males, 46.39g vs 42.71g, P=0.06; females, 41.58 g vs 37.84 g, P=0.24). We also measured lean mass, fat mass, and body composition in Spin4 mutant and wild-type mice, but did not find Spin4 ablation to have any significant effect on body composition (**Figure 1C-E**). Finally, we compared bone mineral density and bone mineral content between wild-type and Spin4 mutant mice and found no noticeable difference in Spin4^Y/-^ or Spin4^+/-^ mice compared to wild-type littermantes (**Figure 1F,G**). Taken together, these data suggest that loss of Spin4 mutation does not affect lean or fat mass accumulation, or bone mineralization in aged mice.

**Figure 1.**
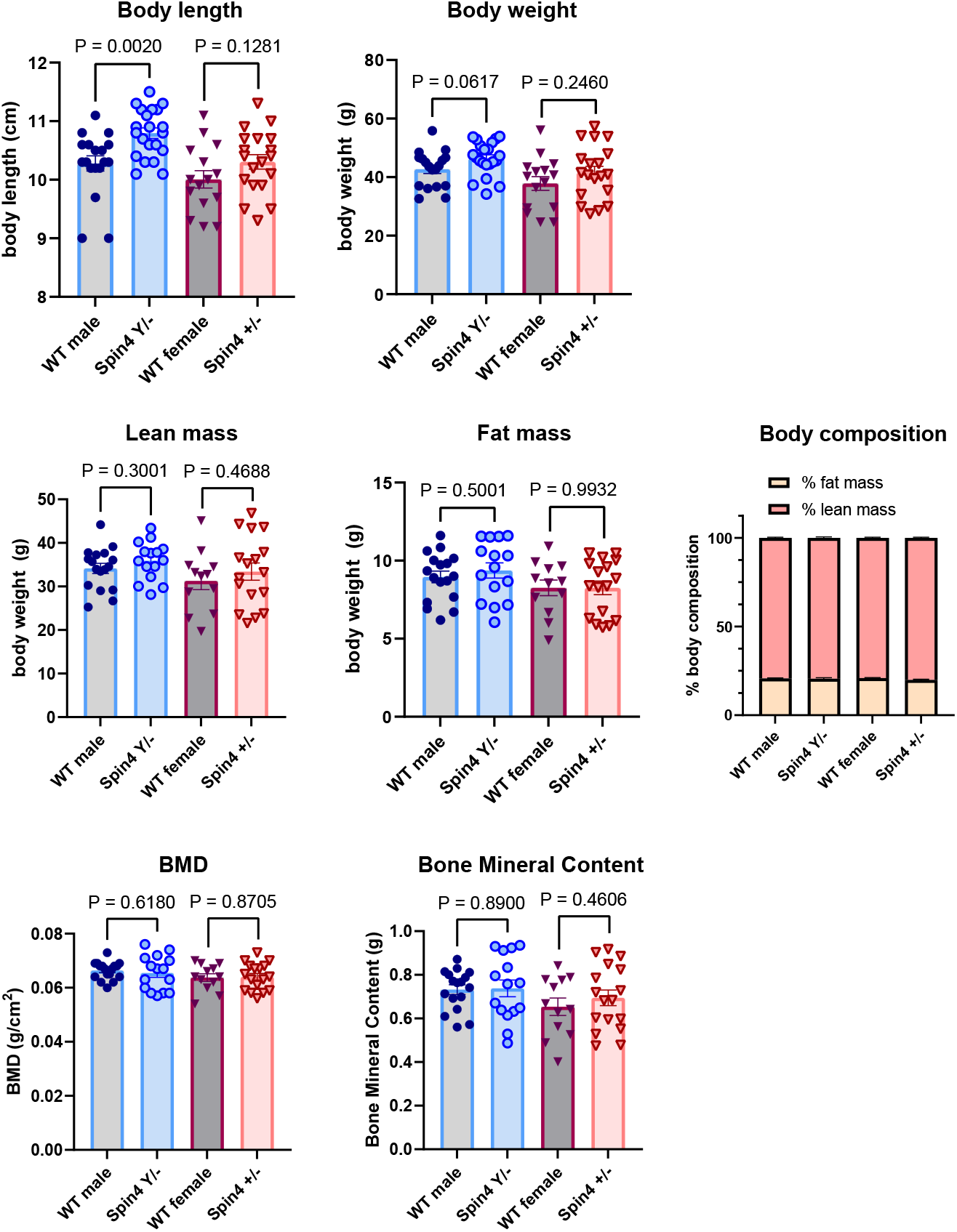
Body size, body composition, and bone density of Spin4 mutant mice and wild-type littermates at 18 months of age. Male and female wild-type (WT, male N=17, female N=12), male hemizygous (Spin4^Y/-^, N=15), or female heterozygous (Spin4^+/-^, N=17) mice were fed ad lib and sacrificed at 18 months of age. Body length was measured from nose to anus. Lean and fat mass, bone mineral density (BMD), and bone mineral content were measured using Xpert 80-L Specimen Radiography System. Bar graphs showing mean ±SEM with individual values in dot plots. P-values represent 2-tailed t-tests, WT vs Spin4 mutant of the same sex.

### Increased neoplasm in Spin4^Y/-^ mice

Mice at 18 months of age were evaluated for tumor formation and/or malignancies by veterinary pathologists. Previous studies have shown evidence that C57BL/6 mice are generally resistant to tumor development, with female being more prone to certain types of cancer (14), such as lymphoma and histiocytic sarcoma (15). In our cohort of 18-month-old wild-type C57/BL6 mice, we identified tumors in 3 out of 32 (9.45%); all were in females (**Table 1**). Two of the tumors were monocytic/histiocytic neoplasms and one was a mesenteric lymphoma. Our findings were consistent with prior literature showing these cancers are more commonly found in aging C57BL/6 female mice than in male mice (15). For the 18-month-old Spin4^Y/-^ (male) or Spin4^+/-^ (female) mice, we found neoplasms in 8 out of 42 mice (19.0%) (**Table 1**). In the female heterozygous Spin4^+/-^ mice, we found three cases of tumors, including one histiocytic sarcoma, one spleen lymphoma, and one mesenteric neoplasm. All these tumor types are not uncommon in aging C57BL/6 female mice (14). There was also no statistical significance difference in prevalence between wild-type and Spin4^+/-^ female mice (3/15 in wild-type and 3/23 in Spin4^+/-^, P=0.66, two-sided Fisher’s exact test). In contrast, we found 5 cases of tumor formation in the male Spin4^Y/-^ mice, including histiocytic sarcoma and mesenteric lymphoma, which are common, but also two cases of bronchiolo-alveolar carcinoma and one cranial osteoma, which are not as common (14) (**Figure 2, Table 1**). The prevalence of tumors found in Spin4^Y/-^ male mice was significantly greater than the prevalence in wild-type littermates (0/17 in wild-type and 5/19 in Spin4^+/-^; P=0.047, two-sided Fisher’s exact test). Our findings therefore support the hypothesis that loss-of-function of Spin4, which is associated with overgrowth in humans and mice, may lead to increased cancer risk.

**Table 1.**
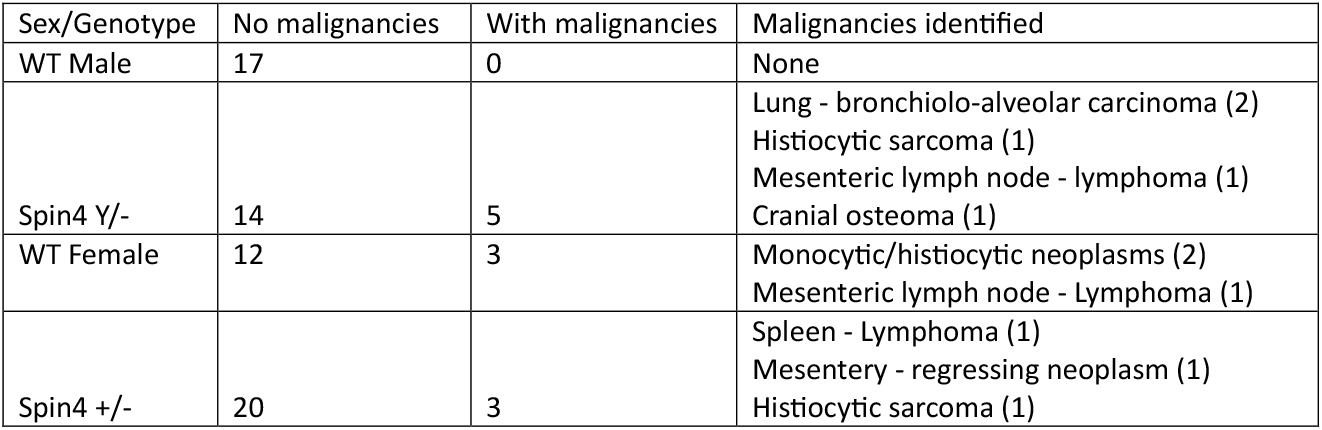
Neoplasms identified in 18-month old wild-type and Spin4 mutant mice.

| Sex/Genotype | No malignancies | With malignancies | Malignancies identified |
| --- | --- | --- | --- |
| WT Male | 17 | 0 | None |
| Spin4 Y/- | 14 | 5 | Lung - bronchiolo-alveolar carcinoma (2)<br>Histiocytic sarcoma (1)<br>Mesenteric lymph node - lymphoma (1)<br>Cranial osteoma (1) |
| WT Female | 12 | 3 | Monocytic/histiocytic neoplasms (2)<br>Mesenteric lymph node - Lymphoma (1) |
| Spin4 +/- | 20 | 3 | Spleen - Lymphoma (1)<br>Mesentery - regressing neoplasm (1)<br>Histiocytic sarcoma (1) |

**Figure 2.**
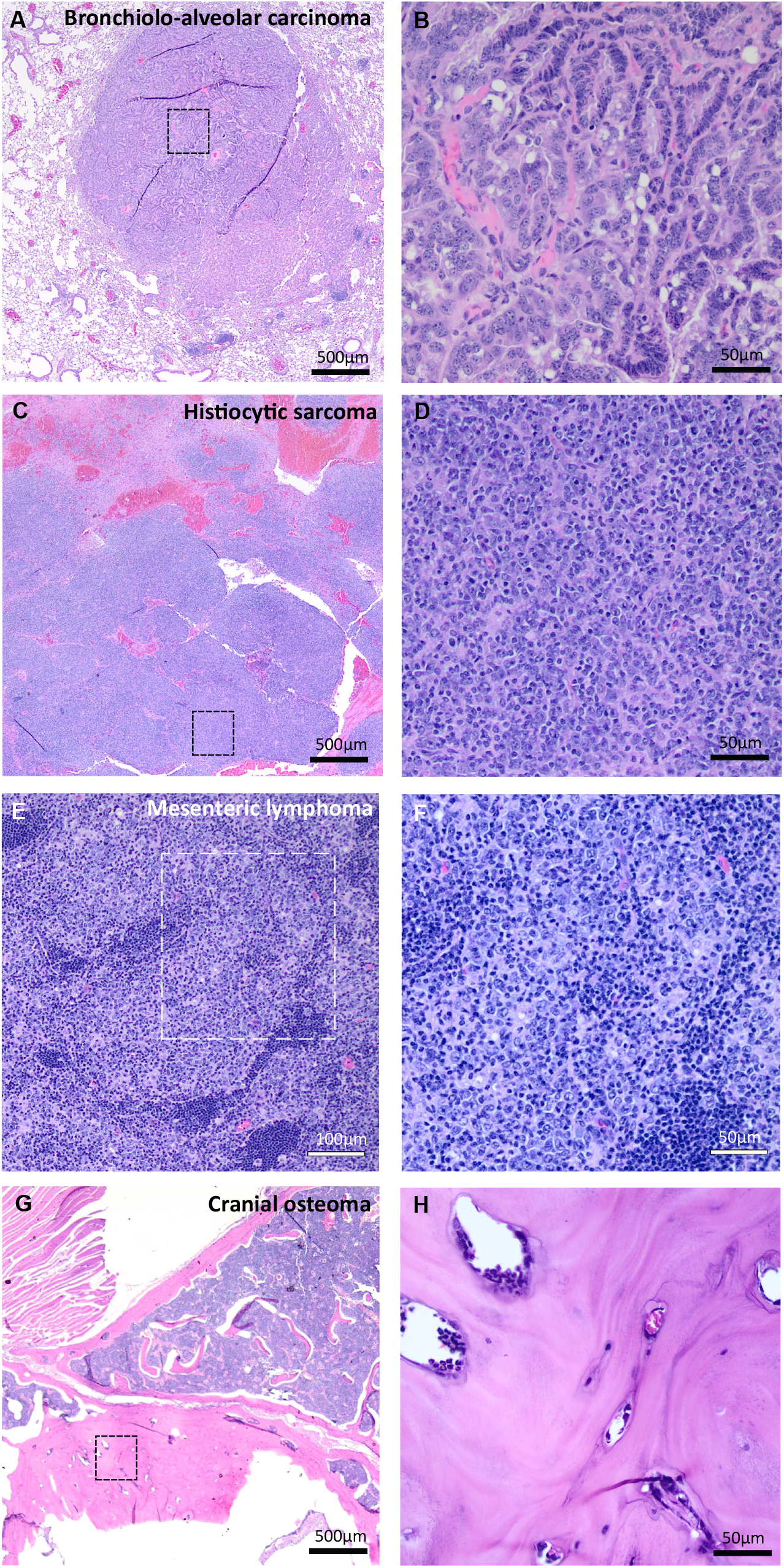
Histopathology of tumors identified in 18-month-old male mice lacking Spin4 (Spin4Y/-). Images on the left are low-magnification images; those on the right are higher-magnification views of the same tumor, taken from the region indicated by the dotted squares. (A,B) Bronchio-alveolar carcinoma found in the lung. The papillae of blue-stained cells represent the malignancy. Large pink cells represent infiltrating macrophage. A similar bronchio-alveolar carcinoma was found in a second Spin4Y/-animal (histology not shown). (C, D) Histiocytic sarcoma found within the mesentery. The pink areas on low magnification represent regions of necrosis and hemorrhage. The dense blue-stained cells represent the malignant histiocytes. (E, F) Lymphoma found within a mesenteric lymph node. The lymphoblasts were found to be invading the adjacent adipose tissue. (G, H) Osteoma found in the cranium. The large pink-staining mass in the center of the figure represents the benign osteoma which consisted of woven bone and was growing inwards from the cranium.

### Bioinformatic analysis of Spin4 expression in human cancer cells

Finally, we compared the level of SPIN4 expression in publicly available RNA-Seq data in 26 human cancer types with their matched normal samples. Based on our findings that Spin4^Y/-^ male mice have increased frequency of tumors in 18-month-old mice, we hypothesized that SPIN4 expression is downregulated in some cancers. Contrary to our prediction, we found that expression of SPIN4 was elevated (P=0.0008) in many human cancers (**Figure 3**). In addition, we performed expression analysis on several other epigenetic regulators that cause overgrowth, including NSD1, EZH2, DNMT3A, and CHD8, and found that, similar to SPIN4, EZH2 (P<0.0001) and DNMT3A (P=0.0057) expression levels were also elevated in many human cancers. As a control, we also analyzed the expression of Cyclin D1, which is often overexpressed in cancers (16), and found that it was elevated (P=0.0002); while expression of Rasgrp2, which is a tumor suppressor (17), was downregulated (P=0.0149), supporting the validity of this analysis.

**Figure 3.**
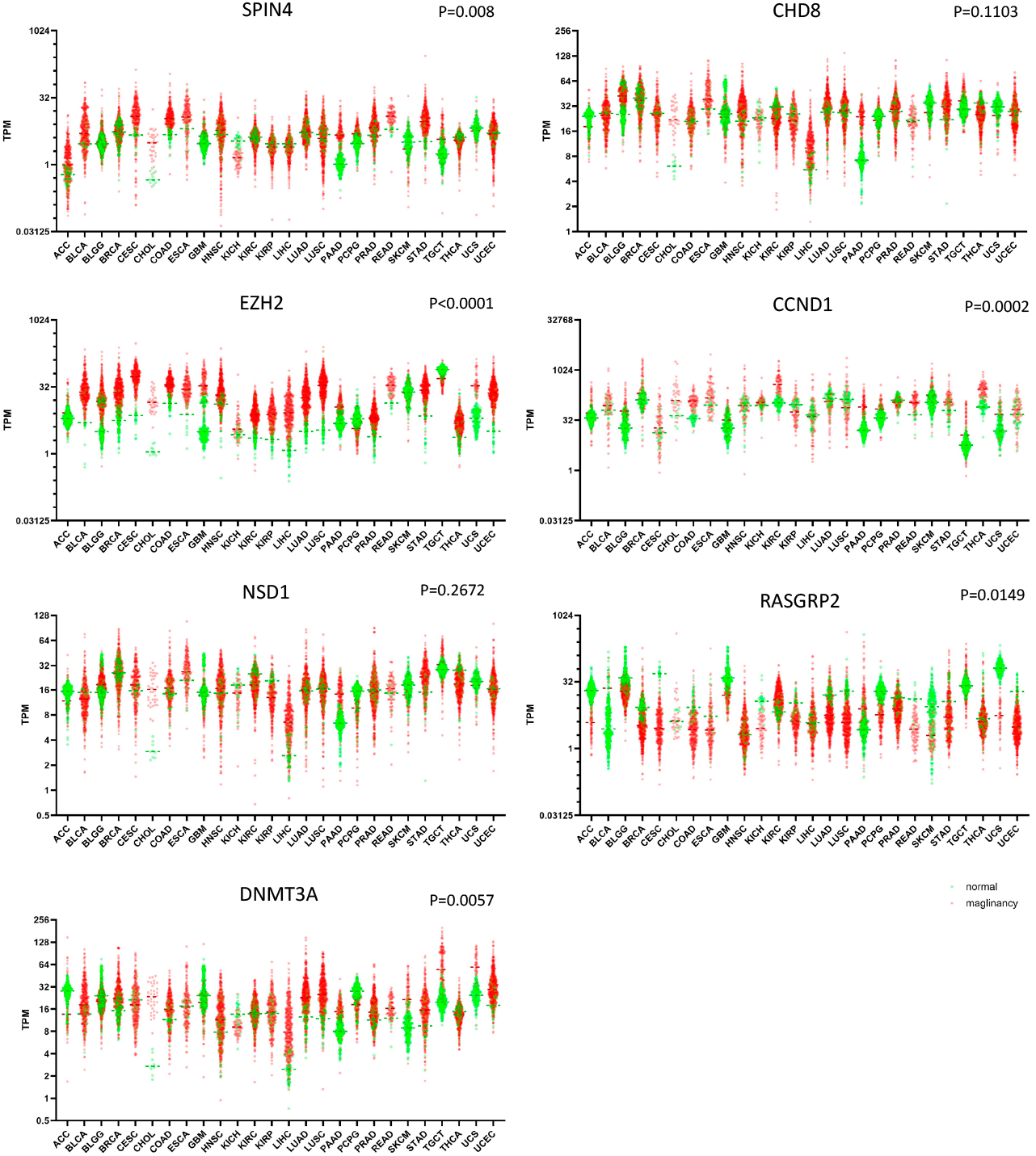
Expression of SPIN4, EZH2, and DNMT3A are elevated in many types of human cancers. Publicly available RNA-Seq data from The Cancer Genome Atlas Program (TCGA) were used to generate scatter plots with each dot representing a single sample. Red, malignancy; green, matched normal sample. P-values represents 2-tailed paired t-tests, normal vs malignancy. CCND1 and RASGRP2 were used as control representing a gene frequently overexpressed in cancer and a tumor suppressor gene, respectively. ACC, adrenocortical carcinoma; BLCA, bladder urothelial carcinoma; BRCA, breast cancer; CESC, cervical squamous cell carcinoma and endocervical adenocarcinoma; CHOL, cholangiocarcinoma; COAD, colon adenocarcinoma; ESCA, esophageal carcinoma; GBM, glioblastoma multiforme; HNSC, head and neck squamous cell carcinoma; KICH, kidney chromophobe; KIRC, kidney renal clear cell carcinoma; KIRP, kidney renal papillary cell carcinoma; LGG, lower grade glioma; LIHC, liver hepatocellular carcinoma; LUAD, lung adenocarcinoma; LUSC, lung squamous cell carcinoma; PAAD, pancreatic adenocarcinoma; PCPG, pheochromocytoma and paraganglioma; PRAD, prostate adenocarcinoma; READ, rectum adenocarcinoma; SKCM, skin cutaneous melanoma; STAD, stomach adenocarcinoma; TGCT, testicular germ cell tumors; THCA, thyroid carcinoma; UCEC, uterine corpus endometrial carcinoma; UCS, uterine carcinosarcoma.

## Discussion

In this study, we compared body length, body weight, body composition, and bone mineral density of 18-month-old Spin4 knockout mice with their wild-type littermates. We found that hemizygous knockout Spin4^Y/-^ male mice have increased body length compared to their male wild-type counterparts. A trend was observed in the heterozygous Spin4^+/-^ female mice but did not reach statistical significance. Similarly, we found a general trend of increased body weight in the Spin4 knockout mice which did not reach statistical significance. These findings are generally consistent with our previous study where we followed body growth of Spin4 mutant mice for up to 10 weeks of age, and found increased body weight, body length, and tibia length in Spin4^Y/-^ male mice (8). Here, we also studied body composition and bone mineral density in these mice at 18 months of age and did not find any significant difference between wild-type and Spin4 mutant mice, suggesting that Spin4 mutation did not significantly affect adipose accumulation, or the balance between bone formation and resorption.

Another important goal of our current study was to investigate whether Spin4 loss leads to an increased risk of neoplasia. We found that, in the hemizygous Spin4^Y/-^ male mice, which lack a functional Spin4 allele, more tumors were detected at 18 months of age, some of which, such as bronchiolo-alveolar carcinoma and cranial osteoma, are not commonly found in wild-type C57BL/6 mice (14). These findings are consistent with the general notion that germline mutations that cause somatic overgrowth tend to be associated with an increased risk of malignancies. Because the P-value in our study was close to the threshold for significance, it will be important to replicate this observation. It would also be important to perform long-term clinical studies in patients with Spin4 overgrowth to evaluate this possible association.

Because we found evidence that loss of Spin4 increased the prevalence of neoplasms in mice, we hypothesized that SPIN4 expression might be decreased in some human malignancies. We therefore analyzed the level of SPIN4 mRNA expression in 26 types of human cancer and their matched normal samples in a publicly-available database and found that, contrary to our hypothesis, SPIN4 expression was elevated in many of the malignancies. However, this observation is consistent with a recent report showing increased SPIN4 expression is associated with advanced nodal status and unfavorable prognosis in nasopharyngeal carcinoma (18). Moreover, analogous findings have been reported for other epigenetic regulators that causes overgrowth. For example, Weaver syndrome, which is caused by a partial loss-of-function mutation of EZH2 (19), is associated with higher risks for several types of malignancies (7) and yet EZH2 is commonly overexpressed in a variety of cancers, such as breast, lung, liver, and ovarian cancers (20), and is classified as an oncogene (21). Similarly, germline loss-of-function mutations in DNMT3A cause human overgrowth in Tatton-Brown-Rahman syndrome while overexpression of DNMT3A has been reported in various cancers (22). Although it remains unclear whether Tatton-Brown-Rahman syndrome is associated with an increased cancer risk, loss-of-function mutations in DNMT3A are frequently found in patients with acute myeloid leukemia (AML) (23), In our current bioinformatic analysis, we found that all three of these genes, SPIN4, EZH2, and DNMT3A tend to be elevated in human cancers compared to normal tissue samples. Based on our observations, we speculate that genes implicated in epigenetic regulation and human overgrowth may require a carefully titrated gene dosage or activity level for normal cell physiology, such that either gain- or loss-of-function would both lead to increased tumorigenesis.

In conclusion, we previously found that loss of SPIN4 causes overgrowth in mice and humans. Here, we studied the long-term effects of Spin4 loss in aged mice. Our findings suggest that loss of this epigenetic reader may predispose to neoplasia but does not affect adiposity or bone density.

## Acknowledgement

This research was supported by the Intramural Research Program of the National Institutes of Health (NIH). The contributions of the NIH author(s) were made as part of their official duties as NIH federal employees, are in compliance with agency policy requirements, and are considered Works of the United States Government. However, the findings and conclusions presented in this paper are those of the author(s) and do not necessarily reflect the views of the NIH or the U.S. Department of Health and Human Services. We thank Dr. Matthew Starost, Dr. Victoria Hoffmann, Dr. Lauren Brinster, and Dr. Michael Eckhaus of the Veterinary Pathology Service, Division of Veterinary Resources, National Institutes of Health (NIH) for providing expert necropsy and histopathology.

## Authors Contribution

JCL and JB conceived the project and designed experiments. JCL, IH, and ARH performed experiments and analyzed data. LD created the mouse model. JCL and JB wrote the manuscript with input from all co-authors.

## Reference

1. Lui JC, and Baron J. Epigenetic Causes of Overgrowth Syndromes. J Clin Endocrinol Metab. 2024;109(2):312–20.

2. Lapunzina P. Risk of tumorigenesis in overgrowth syndromes: a comprehensive review. Am J Med Genet C Semin Med Genet. 2005;137c(1):53–71.

3. Connolly GK, Harris RD, Shumate C, Rednam SP, Canfield MA, Plon SE, et al. Pediatric cancer incidence among individuals with overgrowth syndromes and overgrowth features: A population-based assessment in seven million children. Cancer. 2024;130(3):467–75.

4. Kalish JM, Becktell KD, Bougeard G, Brodeur GM, Diller LR, Doria AS, et al. Update on Surveillance for Wilms Tumor and Hepatoblastoma in Beckwith-Wiedemann Syndrome and Other Predisposition Syndromes. Clin Cancer Res. 2024;30(23):5260–9.

5. Neri G, Gurrieri F, Zanni G, and Lin A. Clinical and molecular aspects of the Simpson-Golabi-Behmel syndrome. Am J Med Genet. 1998;79(4):279–83.

6. Bennett RL, Swaroop A, Troche C, and Licht JD. The Role of Nuclear Receptor-Binding SET Domain Family Histone Lysine Methyltransferases in Cancer. Cold Spring Harb Perspect Med. 2017;7(6).

7. Nakagawa M, and Kitabayashi I. Oncogenic roles of enhancer of zeste homolog 1/2 in hematological malignancies. Cancer Sci. 2018;109(8):2342–8.

8. Lui JC, Wagner J, Zhou E, Dong L, Barnes KM, Jee YH, et al. Loss-of-function variant in SPIN4 causes an X-linked overgrowth syndrome. JCI Insight. 2023;8(9).

9. Chawla N, Rajput M, Shriyan R, Bachani S, and Gupta A. Clinical Observation of a Rare Binder Phenotype With Fetal Overgrowth Due to a SPIN4 Mutation. Matern Fetal Med. 2025;7(4):258–9.

10. Sotos JF, Dodge PR, Muirhead D, Crawford JD, and Talbot NB. CEREBRAL GIGANTISM IN CHILDHOOD. A SYNDROME OF EXCESSIVELY RAPID GROWTH AND ACROMEGALIC FEATURES AND A NONPROGRESSIVE NEUROLOGIC DISORDER. N Engl J Med. 1964;271:109–16.

11. Gibson WT, Hood RL, Zhan SH, Bulman DE, Fejes AP, Moore R, et al. Mutations in EZH2 cause Weaver syndrome. Am J Hum Genet. 2012;90(1):110–8.

12. Tatton-Brown K, Seal S, Ruark E, Harmer J, Ramsay E, Del Vecchio Duarte S, et al. Mutations in the DNA methyltransferase gene DNMT3A cause an overgrowth syndrome with intellectual disability. Nat Genet. 2014;46(4):385–8.

13. O’Roak BJ, Vives L, Fu W, Egertson JD, Stanaway IB, Phelps IG, et al. Multiplex targeted sequencing identifies recurrently mutated genes in autism spectrum disorders. Science. 2012;338(6114):1619–22.

14. Elies L, Guillaume E, Gorieu M, Neves P, and Schorsch F. Historical Control Data of Spontaneous Pathological Findings in C57BL/6J Mice Used in 18-Month Dietary Carcinogenicity Assays. Toxicologic Pathology. 2024;52(2-3):99–113.

15. Nakamura K-i, Kuramoto K, Shibasaki K, Shumiya S, and Ohtsubo K. Age-related Incidence of Spontaneous Tumors in SPF C57BL/6 and BDF<SUB>1</SUB> Mice. Experimental Animals. 1992;41(3):279–85.

16. Montalto FI, and De Amicis F. Cyclin D1 in Cancer: A Molecular Connection for Cell Cycle Control, Adhesion and Invasion in Tumor and Stroma. Cells. 2020;9(12).

17. Liu Y, Ouyang Y, Feng Z, Jiang Z, Ma J, Zhou X, et al. RASGRP2 is a potential immune-related biomarker and regulates mitochondrial-dependent apoptosis in lung adenocarcinoma. Front Immunol. 2023;14:1100231.

18. Chang SL, Chan TC, Chen TJ, Yang CC, Tsai HH, Yeh CF, et al. High SPIN4 Expression Is Linked to Advanced Nodal Status and Inferior Prognosis in Nasopharyngeal Carcinoma Patients. Life (Basel). 2021;11(9).

19. Lui JC, Barnes KM, Dong L, Yue S, Graber E, Rapaport R, et al. Ezh2 Mutations Found in the Weaver Overgrowth Syndrome Cause a Partial Loss of H3K27 Histone Methyltransferase Activity. J Clin Endocrinol Metab. 2018;103(4):1470–8.

20. Eich ML, Athar M, Ferguson JE, 3rd, and Varambally S. EZH2-Targeted Therapies in Cancer: Hype or a Reality. Cancer Res. 2020;80(24):5449–58.

21. Guan X, Deng H, Choi UL, Li Z, Yang Y, Zeng J, et al. EZH2 overexpression dampens tumor-suppressive signals via an EGR1 silencer to drive breast tumorigenesis. Oncogene. 2020;39(48):7127–41.

22. Zhang W, and Xu J. DNA methyltransferases and their roles in tumorigenesis. Biomark Res. 2017;5:1.

23. Ley TJ, Ding L, Walter MJ, McLellan MD, Lamprecht T, Larson DE, et al. <i>DNMT3A</i> Mutations in Acute Myeloid Leukemia. New England Journal of Medicine. 2010;363(25):2424–33.

